# A spatial genomic approach identifies time lags and historic barriers to gene flow in a rapidly fragmenting Appalachian landscape

**DOI:** 10.1101/777920

**Authors:** Thomas A. Maigret, John J. Cox, David W. Weisrock

## Abstract

The resolution offered by genomic data sets coupled with recently developed spatially informed analyses are allowing researchers to quantify population structure at increasingly fine temporal and spatial scales. However, uncertainties regarding data set size and quality thresholds and the time scale at which barriers to gene flow become detectable have limited both empirical research and conservation measures. Here, we used restriction site associated DNA sequencing to generate a large SNP data set for the copperhead snake (*Agkistrodon contortrix*) and address the population genomic impacts of recent and widespread landscape modification across an approximately 1000 km^2^ region of eastern Kentucky. Nonspatial population-based assignment and clustering methods supported little to no population structure. However, using individual-based spatial autocorrelation approaches we found evidence for genetic structuring which closely follows the path of a historic highway which experienced high traffic volumes from ca. 1920 to 1970. We found no similar spatial genomic signatures associated with more recently constructed highways or surface mining activity, though a time lag effect may be responsible for the lack of any emergent spatial genetic patterns. Subsampling of our SNP data set suggested that similar results could be obtained with as few as 250 SNPs, and thresholds for missing data exhibited limited impacts on the spatial patterns we detected outside of very strict or permissive extremes. Our findings highlight the importance of temporal factors in landscape genetics approaches, and suggest the potential advantages of large genomic data sets and fine-scale, spatially-informed approaches for quantifying subtle genetic patterns in temporally complex landscapes.

## Introduction

Habitat loss and fragmentation resulting from natural resource extraction, agriculture, and urbanization is setting some populations on new demographic trajectories, with increasing and persistent genetic diversity loss (Haddad et al. 2015). Understanding the effects of this rapid landscape change on population structure and genetic diversity is critical for informing science-based conservation and management (Hilty et al. 2012, Keller et al. 2015, Waits et al. 2016). However, a variety of geographic and ecological factors can affect the amount and rate at which spatial genetic structuring builds in a given system, creating challenges for the development of proactive management plans (Epps and Keyghobadi 2015, Balkenhol et al. 2016, Richardson et al. 2016). Thus, while migration may be limited by contemporary landscape factors, genetic structure may not be detectable until many generations after a barrier forms, especially if the power to detect such patterns is limited by the quantity or quality of genetic data available (Landguth et al. 2010, McCartney-Melstad et al. 2018).

The use of large single nucleotide polymorphism (SNP) data sets has improved the detection of recent habitat fragmentation in several ways. First, increased genome-wide sampling reduces the number of individuals needed to quantify differentiation among sampling locations (Willing et al. 2012, Nazareno et al. 2017). With this lower threshold for per-locale individual sampling, genomic data can permit sampling schemes encompassing a broader geographic area and a more hierarchical design, thus allowing for more robust resolution of patterns at multiple spatial scales (Anderson et al. 2010, Balkenhol and Fortin 2016). Furthermore, while the relatively high mutation rate of microsatellites is advantageous for detecting recent genetic change (Epps and Keyghobadi 2015), the greater genome-wide sampling of large SNP data sets can potentially detect weaker spatial genetic patterns resulting from relatively recent or porous barriers to gene flow (Landguth et al. 2012). For example, McCartney-Melstad et al. (2018) found that with as few as 300-400 SNPs, genetic structure associated with the barrier effects of roads could be detected in amphibian populations where 12 microsatellite loci had previously indicated no structure. SNP data sets of this size are now readily available through methods such as restriction site-associated DNA sequencing (RADseq), allowing for the generation of thousands of loci from non-model organisms with a range of ecological characteristics that may make them prone to the genetic effects of recent habitat fragmentation (Epps and Keyghobadi 2015).

While traditional methods of testing for spatial genetic patterns, such as model-based clustering (e.g., STRUCTURE, Pritchard et al. 2000) or non-parametric exploratory data analyses (e.g., DAPC, Jombart et al. 2010), have been used to characterize genetic diversity across a given area (François and Waits 2015), other methods which are able to separate spatial and non-spatial genetic variation may be better equipped to detect patterns of genetic differentiation in recently fragmented systems or those with high rates of gene flow (Jombart et al. 2008, Galpern et al. 2014). These methods use spatial autocorrelation to tease apart patterns of inter-versus intra-population genetic variation, improving the identification of population structure at fine geographic scales (Galpern et al. 2012). When coupled with genomic data, spatially-informed analyses may also allow for the detection of weak spatial structure related to recent habitat fragmentation or incomplete barriers to migration (Richardson et al. 2016, Richardson et al. 2017, Combs et al. 2018, Combs et al. 2018a). However, alongside these methodological improvements, work remains to understand the amount of genomic data necessary to assess spatial genetic patterns (e.g., McCartney-Melstad et al. 2018), and the effects of genomic data quality on the resolution of recently evolved population structure.

Identifying spatial genetic patterns associated with landscape features is especially pertinent in regions experiencing rapid and recent landscape change. Few regions have experienced this change as rapidly as central Appalachia in the eastern United States, chiefly as a result of the large-scale surface coal mining practices often referred to as ‘mountaintop removal’ (Wickham et al. 2007, Drummond and Loveland 2010, Pericak et al. 2018, Maigret et al. 2019). Alongside mining, the wholesale construction of several high-traffic road systems in the 1970s and 1980s, in part to facilitate the transportation needs of the mining sector, have further subdivided what was formerly a relatively continuous forest landscape with scant high-traffic roads (KTC 2018). Despite the scale of these changes, the effects on native biodiversity are not well understood (Wickham et al. 2013). Given the historically rugged terrain of Appalachia, topographic homogenization produced by surface mining may facilitate dispersal of terrestrial fauna not encumbered by the radically altered soils, flora, and thermal regimes of reclaimed minelands (Wickham et al. 2013), and highways may also facilitate movement in some species (Trombulak and Fissell 2000). Alternatively, less vagile taxa that rely on sparsely distributed microhabitats may be more sensitive to the effects of forest fragmentation, especially if they are susceptible to road mortality.

We sought to understand the impact of recent and major landscape changes on the population structure of the copperhead (*Agkistrodon contortrix*), an abundant snake in eastern Kentucky (Barbour 1962) generally not capable of long-distance (> 1km) individual movements (Sutton et al. 2017). Copperheads rely on rocky hibernacula located high on steep-sided, south-facing slopes for overwintering (Maigret and Cox 2018), sites disproportionately destroyed by surface mining (Maigret et al. 2019). Copperheads are also generally intolerant of dense, invasive vegetation common to many reclaimed surface mines (Carter et al. 2015, Carter et al. 2017). Additionally, herpetofauna generally, and pit vipers in particular, have been shown to be especially vulnerable to vehicular traffic (Andrews and Gibbons 2005, Shepard et al. 2008), and elevated genetic differentiation associated with highways has been detected using microsatellite markers (Clark et al. 2010, DiLeo et al. 2013). Pertinent to this point, our study area contains several major highways [> 3,000 Annual Average Daily Traffic (AADT)] constructed between 1970 and 1985 which could be barriers to movement for *A. contortrix*. Nearly all these highways were constructed along new paths and do not follow major hydrological or topographic features for the majority of their route through the study area. From at least 1900 until 1970, however, the only highway across the study area was KY State Route 476 (formerly old KY State Route 15; hereafter referred to as KY-476). Prior to 1970, KY-476 was a major thoroughfare through the region, following the sinuous course of Troublesome Creek, a tributary of the North Fork of the Kentucky River. This constrained vehicle speed and made the route prone to frequent flood damage, prompting the development and opening of the new KY-15 c. 1975, leading to markedly decreased traffic volumes on KY-476 (∼500 vehicles/day) and pushing most traffic, including many coal-industry commercial vehicles to the new KY-15 (∼ 5,000 vehicles/day).

Using RADseq data and nonspatial and spatially informed analyses, we investigated the potential for recently formed population structure across *A. contortrix* in eastern Kentucky as a result of this landscape change, with a particular focus on the effects of habitat fragmentation via surface coal mining and through the network of historic (c. 1920) and more recently-constructed (c. 1975) high-traffic roads. Specifically, we aimed to: (1) understand the extent and scale of spatial genomic structuring in copperheads across what was until recently a heavily forested landscape; (2) test for associations between current landcover classes and patterns of spatial genomic diversity; (3) identify current or historic linear landscape features which are associated with reduced gene flow, and (4) understand how the size and quality of a data set can affect our ability to detect spatial genomic patterns. More broadly, we aim to shed light on the temporal scale at which barriers to gene flow are detectable using large SNP data sets, to investigate the potential role of spatially-informed methods for identifying recent or weak genetic boundaries, and to provide a starting point for future research into the spatial genetic implications of increasingly popular methods of surface mining.

## Methods

### 2.1 Sampling methods

We sampled *A. contortrix* individuals from an approximately 1,000 km^2^ area of Breathitt, Knott, and Perry counties in eastern Kentucky, USA (Figure 1). We used a hierarchical sampling strategy, sampling at arrays of four to six individual capture sites (Figure 1A). Each individual capture site consisted of a location where a combination of artificial cover and visual encounter surveys were used to capture snakes. An array was composed of at least four individual capture sites, separated by 2-3 km, and arranged roughly in a cross or an ‘x’. In turn, each array was separated by roughly 10-20 km, providing comparisons at multiple spatial scales both within and between each array (Balkenhol and Fortin 2016). Between May 2014 and September 2016, individuals were captured at sampling sites, typically under artificial cover (e.g., sheet metal or other debris). We augmented this design by including individuals captured apart from designated individual capture sites; typically, these snakes were found alive or dead on roadways within the study area or were killed and/or donated by area residents who were able to provide precise locality information for each tissue sample. Live-captured snakes were restrained and two ventral scales were removed, placed in 95% ethanol, and subsequently frozen at −80°C (Maigret 2019). Muscular tissue from the tails of dead snakes was treated similarly.

**Figure 1:**
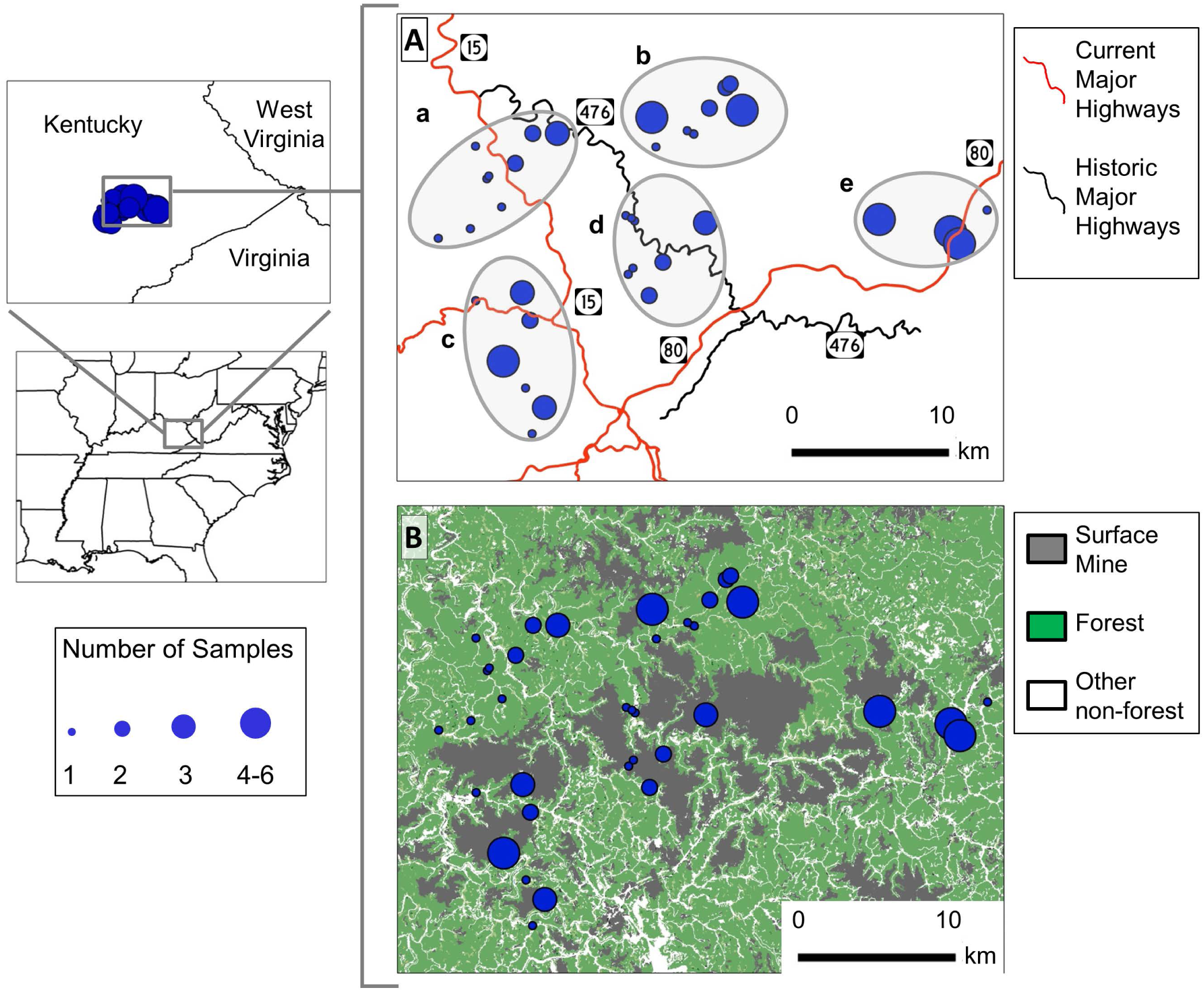
Map of our study area and sampling localities superimposed over (A) the current and historic highway network, with sampling arrays (indicated with a-e) to which each sampling site belongs, and (B) landcover, including surface coal mines, forest, and other non-forest habitat.

### 2.2. DNA sequencing and SNP calling

We extracted genomic DNA using a Qiagen DNeasy Blood and Tissue Extraction Kit and prepared double digest RADseq (ddRADseq) libraries based on Peterson et al. (2012). DNA was quantified using a Qubit 2.0 flourometer (Thermo-Fisher). DNA extractions ≤ 2.0 ng/uL were amplified using a Qiagen REPLI-g high-fidelity whole genome amplification kit. We prepared ddRADseq libraries using ∼1000 ng of DNA per individual. DNA was digested using EcoRI and SphI and subsequently cleaned with Agencourt Ampure XP beads (Beckman Coulter). Adaptor ligation was performed using one of 48 unique 5 bp barcodes in combination with a universal 6 bp single-index PCR adaptor. Samples were then pooled in groups of 8, bead cleaned, and size selected (526 bp ± 10%) with a Pippin Prep (Sage Science). Each 8-sample pool was then Qbit quantified and amplified using Phusion high-fidelity PCR (New England Biolabs) with a PCR primer with one of three unique barcodes, permitting each individual to be identified uniquely using a combination of the unique PCR barcode and a unique adaptor index. After cleaning and quantifying PCR product, we used an Agilent 2100 Bioanalyzer to confirm target fragment size distributions before 150 bp paired-end sequencing on two lanes of an Illumina Hi-Seq 2500. Individuals were randomly assigned to a lane with respect to geographic location to reduce downstream genetic artefacts (Meirmans 2015).

We used Stacks v1.37 (Catchen et al. 2013) to identify orthologous loci across individuals. No overlap was expected between sequencing reads; therefore, we used a custom script to stitch together forward and reverse reads. We used process_radtags to demultiplex individuals and discard low-quality reads containing uncalled bases or a mean quality score < 10 in a sliding window comprising 15% of the read. After quality filtering, reads were assembled using denovo_map, with a minimum stack depth of 5 (m = 5), 3 mismatches allowed between stacks within individuals (M = 3), and 2 mismatches allowed between stacks among individuals (n = 2). To increase confidence in our SNP calls, we used rxstacks to remove SNPs with a low log likelihood (--ln_lim = −25) and/or a high proportion of confounded loci (conf_lim = 0.25). After running rxstacks, cstacks and sstacks were re-run with the filtered loci. We sampled a single SNP per locus (--write-single-snp), using only SNPs with < 50% missing data, a minor allele frequency < 0.015, and no evidence of excess heterozygosity. Finally, we removed individuals with > 50% missing data.

### 2.3 Summary statistics and distance-based analyses

We generated a genetic dissimilarity matrix using the program bed2diffs v1 in the EEMS package (Petkova et al. 2016). This produced a matrix of average individual pairwise genetic dissimilarity (hereafter referred to as the “GDM”) based on allelic frequencies, similar to the proportion of shared alleles (Bowcock et al. 1994). We estimated effective population size using the linkage disequilibrium method in NeEstimator (Do et al. 2014). We estimated heterozygosity and nucleotide diversity using plink and vcftools, respectively (Purcell et al. 2007, Danecek et al. 2011). We calculated the relationship between geography and the GDM using the ecodist package in R. Mantel correlograms were generated for multiple geographic distances, including Euclidean distance, the natural log of distance, stream (hydrological) distance, and the natural log of stream distance. To quantify stream distances, we used the Origin-Destination Cost Matrix tool in ArcMap v10.1 (ESRI, Redlands, CA) and a shapefile of USGS stream paths obtained from the KY Division of Geographic Information. We chose to use stream paths as a proxy for potential elevation effects given the rugged terrain of the study area.

### 2.4 Nonspatial analyses of population structure

To identify and characterize genetic clusters across our study area, we used both discriminant analysis of principal components (DAPC) in the adegenet R package (Jombart et al. 2010) and Bayesian clustering via STRUCTURE (Pritchard et al. 2000). For our DAPC analyses, we first used the find.clusters function, retaining all principal components (PCs) and selecting the *K* value with the lowest Bayesian information criterion (BIC). Individuals were then ordinated in PC space using the dapc function. To reduce the potential for over-fitting, we selected the number of retained PCs in light of diminishing returns from retaining excess PCs (Jombart et al. 2010).

For our STRUCTURE analyses, we estimated population assignment of individuals using an admixture model with cluster numbers ranging from *K* = 1 to 10. Five replicates were run for each *K*, each for 1,000,000 generations after a burn-in of 100,000 generations. We used Structure Harvester v0.6.9.4 (Earl and von Holt 2012) to generate mean log likelihood values for each *K* and identify the optimal number of clusters for our data using δ*K* (Evanno et al. 2005). We used the program CLUMPAK to compute cluster membership coefficients across replicates (Kopelman et al. 2015).

### 2.5 Spatially informed analyses of population structure

To further test for genetic structure across our study system, we used three recently developed approaches that integrate spatial information into analyses based on genetic dissimilarity. First, we used MEMGENE (Galpern et al. 2014), a regression-based analysis based on the spatial autocorrelation among a given set of georeferenced individuals and a corresponding GDM. Individual samples are mapped based on geographic location, and significant eigenvector scores are overlaid to provide a visualization of the geographic nature of genetic dissimilarity among individuals.

Second, we used the program sPCA (Jombart et al. 2008) implemented in the R package adegenet. sPCA is broadly similar to MEMGENE (but see Galpern et al. 2014:Appendix S4), but relies on an ordination approach based on Moran’s *I* index to identify eigenvectors which maximize variation in allele frequencies and spatial autocorrelation, and then maps these eigenvectors on to geographic coordinates. Our analyses used a nearest-neighbor connection network with k = 40 neighbors to maximize potential connectivity across our large number of spatially distinct samples. We relied on the eigenvalue variance and spatial components plots to select the optimal number of global and local axes to retain, and used the recommended multivariate significance test to identify significant global and local genomic structure.

Third, to take into consideration the impact of landscape features on gene flow, we estimated a resistance model using the R package ResistanceGA (Peterman 2018). ResistanceGA uses a genetic algorithm approach to optimize the individual resistance values associated with a given resistance surface based on genetic dissimilarity data. Model fit of the optimized surfaces are quantified using AIC values from linear mixed-effects models, both for each surface individually and for all combinations of individual surfaces. Thus, ResistanceGA bypasses the often-subjective expert opinion parameterization stage of resistance surface construction (Spear et al. 2016, Peterman et al. 2018). Our ResistanceGA input landscape surfaces consisted of land cover classification data obtained from 2011 National Land Cover Data. We reclassified raw NLCD raster values into 3 different resistance surfaces of two categories each, including: (1) a mining surface with two categories, mined and unmined land, (2) a surface representing the route of current highways, with two categories, highway and non-highway, and (3) a surface representing the route of KY-476, also with two categories, highway and non-highway (Table S1). We tested both for effects of each of these three surfaces independently and each possible combination of the three. We reclassified NLCD raster classes using the Reclassify tool in the Spatial Analyst extension of ArcMap 10.3.3, producing our three putative resistance surfaces. We relied on historic road maps publicly available from the KY Transportation Cabinet to identify current and historic highway patterns in the study area from 1936 to the present, and historic topographic maps from the US Geological Survey’s Historical Topographic Map Explorer for information on routes before 1936. Our response data set was our individual pairwise GDM, which we ran alongside our land cover raster surface using the ‘costDistance’ function in the R package gdistance (van Etten 2017). This function calculates least cost paths between each pair of locations, and while lacking the comprehensive approach available with random walk commute times, least cost paths represent a much more computationally tractable approach for our spatial and genetic data set (Peterman 2018).

### 2.6 Subsampling of our SNP data set

We aimed to assess the resolving power of two of the spatially informed methods, sPCA and MEMGENE, based on: (1) the number of loci, and (2) the amount of missing data. To examine the effect of the number of loci, we randomly subsampled our full 2,140 SNP data set, producing subsets of 25, 50, 100, 250, 500, and 1,000 loci. For each subset, ten replicates were generated using plink and analyzed in sPCA and MEMGENE as described above for the full data set. To examine the effects of missing data, we re-filtered our raw SNP data set using missing data thresholds of 0.05 (i.e., retaining only loci present in ≥ 95% of individuals), 0.10, 0.25, 0.40, 0.5, 0.75, 0.90, and 0.95. While these represented our thresholds, our realized data sets typically had smaller amounts of missing data, in aggregate, than each threshold. Only a single data set could be produced for each missing data threshold.

Differences in the how sPCA and MEMGENE are designed influenced how we quantified our subsampling and missing data threshold results. For sPCA, we first detected significant patterns of structuring, then tabulated the proportion of replicates with unrelated, similar, or identical spatial genomic patterns as detected in analysis of the full data set. Model outputs for sPCA include global and local permutation tests of structuring, the p-values of which were obtained for each level of subsampling and missing data thresholds, which we averaged across ten replicates for the former. MEMGENE, on the other hand, only analyzes significant spatial patterns, and nonsignificant patterns are not retained for downstream analyses. Thus, for MEMGENE, we obtained R^2^ values only for levels of subsampling and missing data thresholds where significant spatial patterns were observed, and we quantified spatial patterns which were unrelated, similar, or identical, in a similar fashion to our sPCA results. We visualized these results by charting p-values from local and global tests from sPCA alongside R^2^ values from MEMGENE for each missing data threshold, and by charting both these statistical values and the proportion of similar patterns for each subsampling level. While categorizing spatial patterns in terms of their similarity to those generated using our full data set requires some qualitative assessment of the results, we chose not to use more substantial quantitative metrics for comparison (e.g., spatial point pattern analysis) given the limited number of sampling sites.

## Results

### 3.1 Sequencing results

We generated ∼239 million 150 bp paired-end reads, with a mean of 1,869,394 reads per individual. Increasing or decreasing the minimum read depth between 4 and 7 did not affect any summary statistics, and only marginally affected the number of loci in our data (Table S2). After filtering, we recovered genotypes for 77 individuals from 34 different locations (Figure 1). This included a total of 2,140 loci, with an average missing data rate of 23.5% of loci per individual (min. = 4.4%; max.= 48.9%).

### 3.2 Summary of genetic diversity

Across our study area, we estimated HO = 0.193, HE = 0.24, π = 0.242, and FIS = 0.195. We estimated an Ne of 635.8 (95% CI: 595.6, 681.6). We identified weak, but sometimes significant correlations between genetic and different measures of geographic distances, including Euclidean (p = 0.79, R^2^ = −0.0003), natural log of Euclidean (p = 0.0006, R^2^ =0.0037), hydrological distance (p < 0.0001, R^2^ = 0.0054), and natural log of hydrological distance (p = 0.003, R^2^ = 0.0029). Mantel correlograms of correlation by distance class similarly showed minimal evidence of isolation-by-distance (Figure S1).

### 3.3 Non-spatial population structure

Neither DAPC nor STRUCTURE analyses supported the presence of multiple geographically distinct genetic clusters. BIC scores in DAPC were lowest for *K* = 1 (Figure 2a), and an exploration of cluster assignments using the first PC axis and a *K* = 2 did not produce individual assignments corresponding to sampling localities or geography. STRUCTURE analyses identified a *K* = 3 as the best-fit clustering model for our data based on the δ*K* statistic (Figure 2b). However, at this level of clustering all individuals were nearly equally assigned to all three clusters, indicating a lack of population structure; these results were similar at a *K* = 2.

**Figure 2:**
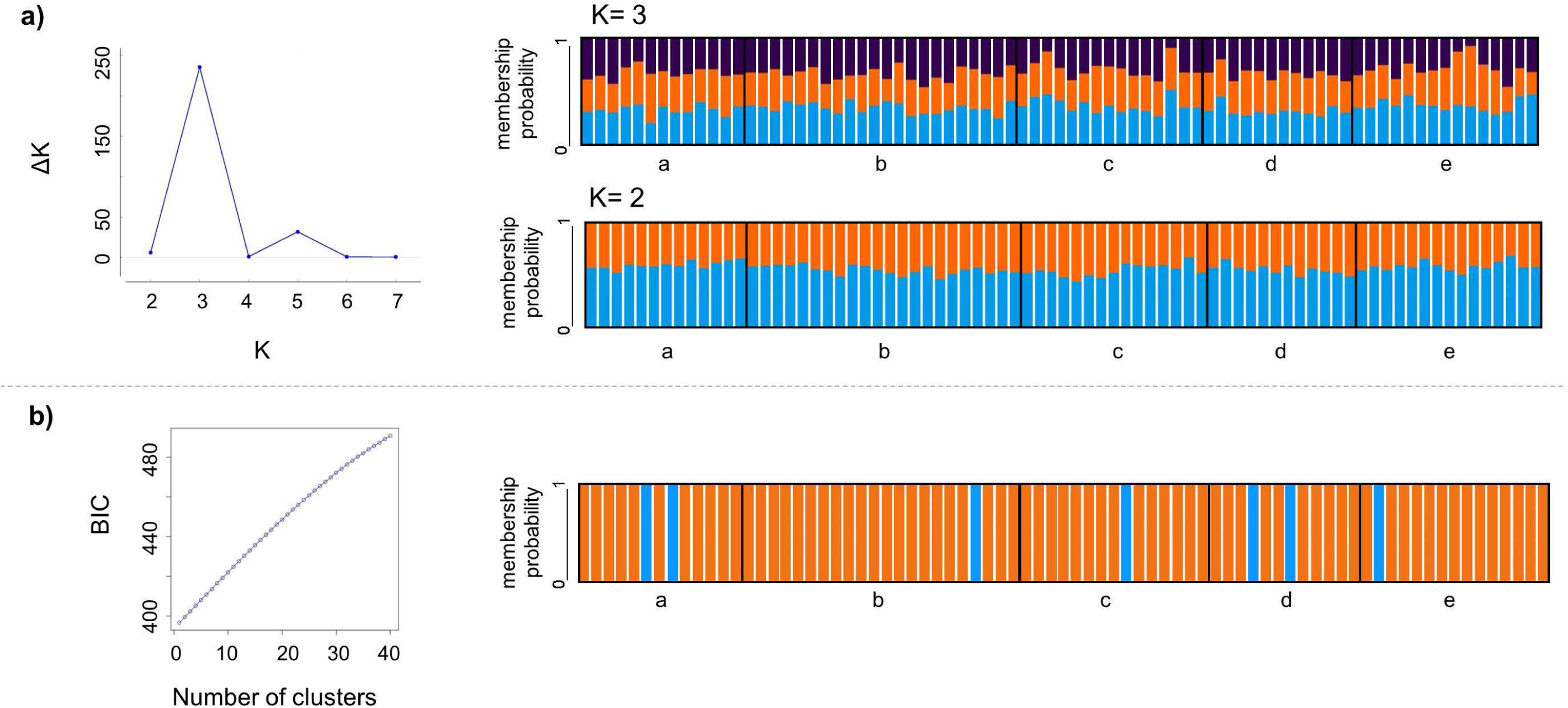
Results of nonspatial population structure analyses, including (a) δ*K* and individual assignment plots from Structure for *K* = 3 and *K* = 2, and (b) BIC and individual assignment plots from DAPC. Letters beneath each individual assignment plot correspond to the geographically distinct sampling arrays depicted in Figure 1a.

### 3.3 Spatially informed population structure

sPCA analyses identified significant global structure across the study area (p = 0.002). The first global (positive) sPCA axis identified a population genetic break that closely followed the path of KY-476 (Figure 3a). Based on a scree plot and a plot of eigenvalues, this first global axis contained the most information relative to other axes, and support for any of the local (negative) axes was not of congruent strength (Figure S2a-b).

**Figure 3:**
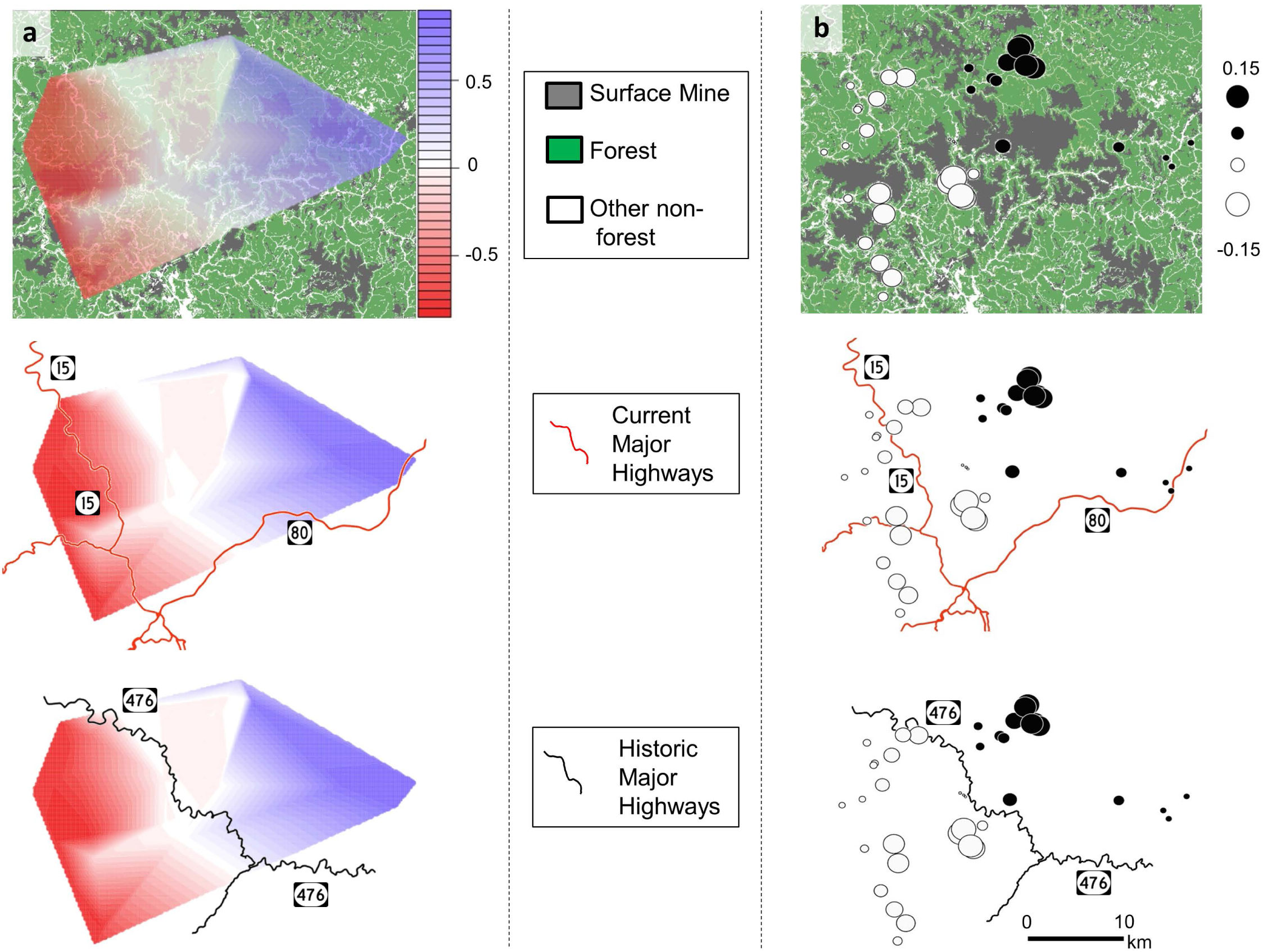
Results of our spatially informed population structure analyses. (a) Results of sPCA analyses visualized using interpolated vector scores, showing divergence coinciding with the historic highway path (designated in this study as KY-476), but not with landcover or current highway infrastructure. (b) Results of MEMGENE analysis, which suggests similar patterns of population structure associated with KY-476. Circle color and size represent the association and genetic similarity, respectively, along the first MEM variable axis.

The first variable identified as significant in the MEMGENE analysis explained a high proportion of the total variance across three retained axes (0.81). The proportion of overall genetic variance explained by spatial patterns associated with this first variable was modest (adj. R^2^ = 0.061), yet was similar to the proportion explained by other studies at similar spatial scales (Galpern et al. 2014, Combs et al. 2018a). Visualization of the first and most explanatory MEM variable similarly identified a genetic break that partitioned populations on either side of KY-476 (Figure 3b). No genetic breaks identified an influence of landcover or current highway paths.

Landscape resistance analyses in ResistanceGA supported a null model of no geographic structure, followed by a model of isolation by distance (Tale S3). Models that included the three individual resistance surfaces (landcover, current highways, or historic highways), or any combination of resistance surfaces, were not strongly supported.

### 3.4 Subsampling of SNP data set

sPCA analysis of subsampled SNP data sets produced significant detection of global structure with as few as 25 loci (average global p-value of ten replicates = 0.067, Fig. 4a), although data sizes ≥ 250 loci were needed to produce identical patterns to those generated with the full data set (mean global p = 0.0033). At ≥ 500 loci identical patterns were produced in all replicates. Significant local structure was not supported for any level of subsampling (mean local p-value = 0.34). MEMGENE analysis of subsampled SNP data sets produced identical patterns in a majority of replicates when sampling ≥ 100 loci (Fig. 4b). However, identical results were still detected in 50% of replicates when sampling 50 loci and produced in all replicates when sampling 1000 loci.

**Figure 4:**
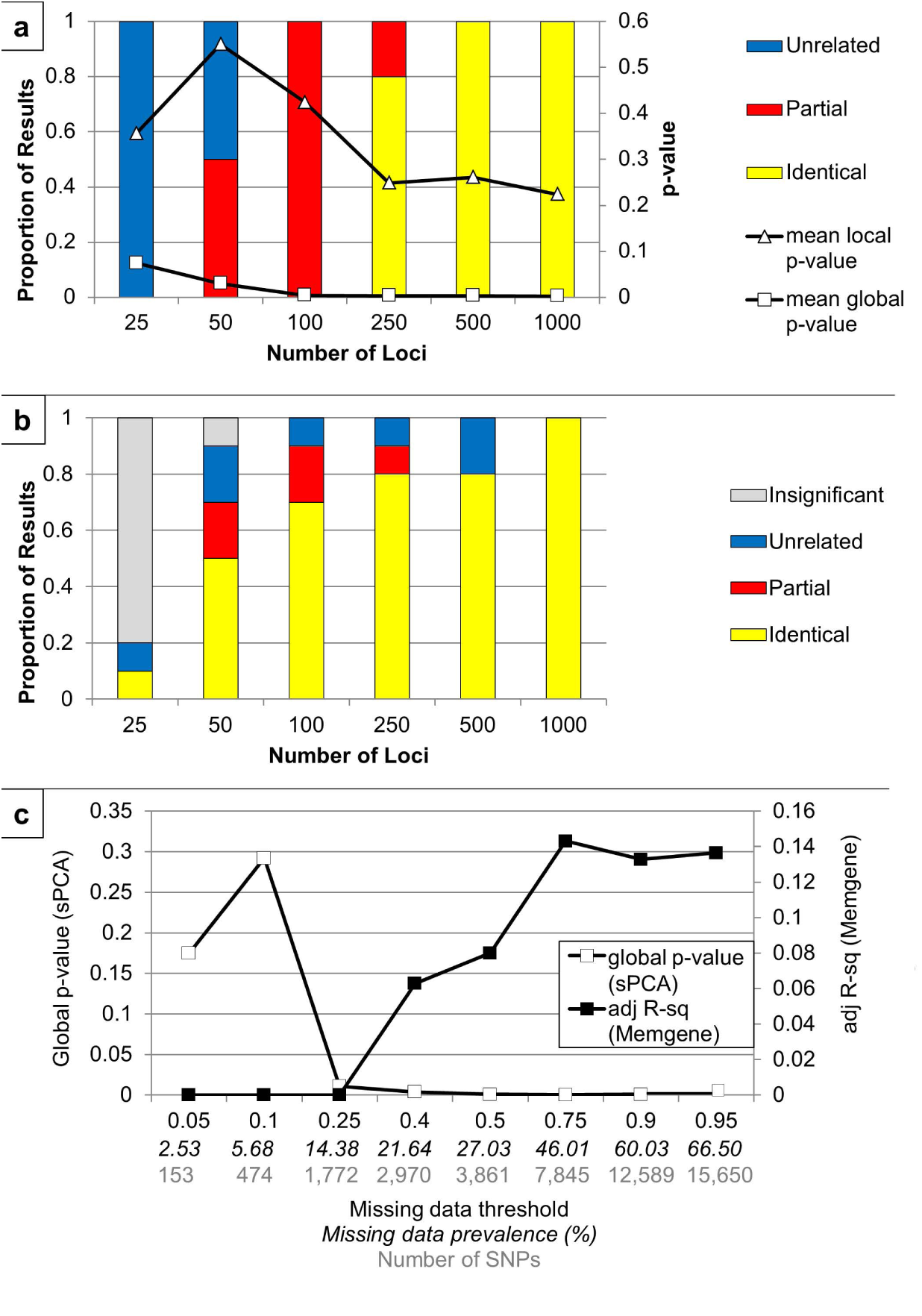
Effects of the number of loci and missing data on (a) sPCA and (b) MEMGENE results, and the effects of missing data levels on both sets of results (c). For (a) and (b), the left y-axis represents the proportion of results which were identical, partially identical, or unrelated to the results obtained from the full data set depicted in Figure 3. Visualization results from each replicate are available in Figure S4. The right y-axis for (a) represents p-values from global and local tests for structuring.

The performance of sPCA and MEMGENE was not adversely affected by the inclusion of higher levels of missing data (Fig. 4c). Global p-values from sPCA analysis were significant at ≥ 25% missing data and remained so, even when the data set allowed for as much as 95% missing data per individual. Missing data levels ≤ 10% resulted in a loss of significant global spatial structure, and we note the peculiarity that data sets of smaller locus number resulted in significant detection of global structure in our subsampled data replicates, suggesting the potential for Type I error with small data sets. Non-zero R^2^ estimates were produced in MEMGENE analyses of data sets permitting ≥ 40% missing and increased with higher levels of missing data, up to 75%.

## Discussion

### Non-spatial vs. spatially informed inference of population structure

Here, we present empirical evidence for the ability of some spatially informed methods to detect weak population structure in study systems where more traditional and non-spatially informed methods indicate a lack of structure. Patterns in both DAPC and Structure results were consistent with a *K* = 1 model, with no evidence for geographically distinct genetic clusters across the study area. In contrast, the spatially informed methods sPCA and MEMGENE returned similar results supporting geographic genetic structure with a break coinciding with the path of KY-476, a historically-important highway that served as a major traffic artery in the region between c. 1920-1975. The inference of weak population structure and genetic fragmentation on the landscape of our study system is bolstered by multiple lines of evidence. First, both sPCA and MEMGENE identified the same geographic genetic break. While these methods both use spatial autocorrelation in the analysis of genetic data, they operate in very different ways: sPCA relies on the integration of Moran’s I matrix via a connection network, while MEMGENE uses a forward selection method to identify significant MEM eigenvectors, and then uses a regression approach to generate output variables which contain the spatial patterns (Galpern et al. 2014). The congruence of these results indicates that our result is probably not a spurious pattern driven by an artefact of one particular analysis. Second, while the magnitude of population structure detected in our work was generally weak—MEMGENE-based regression analyses attributed ∼6% of total genetic variation to spatial effects—this amount of spatially explained genetic variation is in the range of that detected with MEMGENE under simulated models of population fragmentation and higher than that detected for panmictic populations (Galpern et al. 2014). This level of spatially driven genetic variation is also similar to that detected in other studies of recently fragmented landscapes (Combs et al. 2018a, Combs et al. 2018b). Our overall interpretation of these results is that the use of methods that specifically use spatial patterns of variation, such as sPCA and MEMGENE, seem to be able to identify patterns of weak population structure at temporal and spatial scales where more widely used non-spatial methods would fail to discern geographic population structure (Galpern et al. 2014).

In contrast, our estimates of population structure using optimized landscape resistance generated using ResistanceGA did not support a link between genetic differentiation and landscape features. This lack of spatially informed population structure may be related to methodological aspects of this program, as it does not use the autocorrelation approach that is built in to sPCA and MEMGENE. In addition, our analyses were limited to only analyzing least cost paths. ResistanceGA does allow for a more exhaustive exploration of restricted gene flow across the landscape using a random-walk framework, which may have identified fragmentation associated with landscape features that were not examined using a least cost path approach. However, this required a computationally prohibitive set of analyses given our level of locality sampling and landcover data resolution. Coupled with the relatively weak nature of the spatial genomic signal associated with the route of KY-476, our ability to detect resistance to gene flow based on landcover classes may have been comparatively limited.

### Data size and quality in the detection of weak population structure

While our large genomic data set may have also increased our ability to detect subtle spatial patterns, random subsampling of our data indicated that thousands of SNPs may not be necessary to detect weak population structure similar to that found with our full data set. In fact, we found that several hundred SNPs may be sufficient to consistently identify weak spatial structure. This result is similar to that of a recent study (McCartney-Melstad et al. 2018), which showed that the use of a more limited set of independent SNPs (∼300-400) was sufficient to recover fine-scale population structure using the non-spatial method Admixture (Alexander et al. 2009) with results similar to those obtained with a larger, more-complete data set (3095 SNPs). Our subsampling work extends this finding, indicating that spatially informed methods of population structure may be equally efficient with relatively modest sized data sets (∼250-500 loci). We do note that minimum locus thresholds will vary based on the intensity of the spatial genetic signal, the number of individuals sampled, and a variety of other factors. However, these developing empirical findings provide an optimistic outlook on the minimum data size required for the detection of weak landscape-level fragmentation.

Our exploration of the inclusion of missing data yielded similarly optimistic results, where, under a wide range of thresholds, missing genotypes did not substantially alter our spatial landscape genomic findings. Using stringent missing genotype thresholds, which also lowered the number of SNPs in the data, actually decreased the spatial signal. Conversely, allowing for more missing data increased the signal of population structure in our data, with a plateau in the level of significance (sPCA) and amount of spatial variation explained (MEMGENE). The effect of missing data in population and evolutionary studies has seen mixed results. Simulation-based results have indicated that missing genotypes in RADseq data can result in substantial biases in a range of population genetic summary statistics, including FST (Arnold et al. 2013). In contrast, the use of more liberal missing data thresholds in RADseq-based phylogenetic studies has provided opportunities to recover phylogenetic patterns not detected using more stringent thresholds (Wagner et al. 2013, Eaton et al. 2017; but see Leaché et al. 2015). This may be due to a bias whereby loci with higher mutation rates, but likely to contain population or phylogenetic information, are eliminated by stringent missing thresholds (Huang and Knowles 2014). The effect of missing data in landscape genomic studies has yet to be thoroughly explored, and we suggest based on our results that some spatially informed analyses may be robust to the recovery of patterns of weak population structure despite the inclusion of a high level of missing genotypes, but that parameter estimation at this geographic scale (e.g., migration rates) may be more strongly influenced. Therefore, when possible, we second the recommendation of others (Wagner et al. 2013, O’Leary et al. 2018) for researchers to explore the sensitivity of their results across a range of different missing data thresholds.

### Copperhead landscape genomics and temporal considerations

Our results further emphasize an association between high-traffic roads and genetic differentiation in pit vipers (Clark et al. 2010, DiLeo et al. 2010, DiLeo et al. 2013, Bushar et al. 2015, Herrmann et al. 2017, but see Weyer et al. 2014). These findings are in addition to field studies that have suggested the outsized role played by road mortality in snakes, and herpetofauna more generally (Andrews and Gibbons 2005, Row et al. 2007, Shepard et al. 2008). Furthermore, our results suggest that the effects of high-traffic roads and associated intense human activity might persist for decades after traffic volumes decline, in line with predictions from simulations (Landguth et al. 2010).

We did not find evidence for a strong influence of surface coal mining on genetic connectivity, which was surprising given the widespread nature of surface mining in the study area, and the wholesale shifts in vegetation, soils, topography, and fauna that characterize the mining and mine reclamation process. Surface mining of coal in Appalachia has a high degree of spatial and temporal variance; portions of mines can exist in various states of reclamation from barren rock to early successional forest, and mining activity can cease for months to years as a result of fluctuating coal prices or labor disputes, thus providing opportunities for animals to maintain genetic connectivity in these novel landscapes. Regardless, we recommend further research into this generally understudied area, as the large scale and radical impacts of this mining practice may well result in detectable impacts in populations of other taxa (Wickham et al. 2013). This may be especially true for species with shorter generation times, smaller population sizes, and more exclusive associations with ridgetop forests (Epps and Keyghobadi 2015, Maigret et al. 2019).

We note that the connection between the identified genetic fragmentation and the historic highway KY-476 is a largely qualitative assessment, and several specific caveats deserve mention. The route of KY-476 corresponds not only to a highway path, but also to a swath of comparatively higher historic human population density and also to the route of Troublesome Creek, either of which could be factors more important than the highway itself. While modeling relative contributions of population density and road mortality is beyond the scope of our study, in terms of parallel geomorphology and hydrology, the historic highway path does not correspond to any major feature which might be expected to seriously reduce movement of copperheads (Figure S3). Other waterways which divide our sampling locations, including Lost Creek and Buckhorn Creek, are of similar size to Troublesome Creek. Moreover, copperheads and other pit vipers regularly cross bodies of water (T. Maigret, unpublished data; Clark et al. 2010), and studies have found that even hydropower reservoirs are ineffective barriers to gene flow in copperheads and similar species (Oyler-McCance and Parker 2010, Levine et al. 2016). More generally, a second caveat is that while we intended our sampling to be hierarchical in design, the broad scales at which genomic patterns exist in our study area means that we are examining a single functional landscape. When possible, landscape-scale replication would provide a more robust assessment of the effects of current and historic landscape features on gene flow in *A. contortrix* and similar taxa (Short Bull et al. 2011). Moreover, assuming we have detected a spatial genomic pattern stemming from historic highway traffic, we have not determined the traffic threshold which would produce a noticeable spatial genetic pattern or the precise time lag which must pass before these patterns become detectable. Other research has suggested that even low amounts of traffic can produce genetic differentiation (Clark et al. 2010), and depending on a variety of demographic characteristics, numerous generations may need to pass before genetic differentiation becomes apparent (Landguth et al. 2010, Epps and Keyghobadi 2015). Thus, while we may have detected the effect of a historic roadway, we have not conclusively ruled out impacts of current roadways, or even low-traffic and unpaved county roads not included in our analysis. In a similar manner, our findings regarding the spatial genetic implications of surface mines should also be understood tentatively.

Our study adds to a growing list highlighting the potential for large SNP data sets to detect weak, recent, or otherwise subtle spatial genomic patterns (Gonzàlez-Serna et al. 2018, McCartney-Melstad et al. 2018, Murphy et al. 2018, Tan et al. 2018). Considering the problems time lags present for conservation planning, the use of large (> 250) SNP data sets and spatially informed analyses of genetic diversity will likely become increasingly important for placing patterns of population structuring in their proper genomic, temporal, and geographic contexts.

## Data Accessibility

Sequence data, SNP calls, landcover rasters, and sample catalogs will be accessible upon acceptance via NCBI’s sequence read archive (SRA) at accession number PRJNA6278371.

## Acknowledgements

Our work was conducted under University of Kentucky Institutional Animal Care and Use Committee (IACUC) Protocol 2012-0954. Funding was provided by a Theodore Roosevelt Memorial Grant (American Museum of Natural History), McIntire Stennis Project KY009031 (US Department of Agriculture, National Institute of Food and Agriculture), a Kerri Casner Environmental Sciences Fellowship (Tracy Farmer Institute for Sustainability and the Environment), and an Ellers-Billing Award (University of Kentucky Appalachian Center). We are grateful for field assistance provided by David Collett, Erwin Williams, Chris Osborne, Ted Sizemore, Neva Williams, Grover Napier, RB Combs, Anthony Campbell, Scotty Brewer, Mark Chaffins, Doran Howard, Taylor Hughes, Rudy Noble, Verle Fugate, the late Geraldine McIntosh, and other residents of Breathitt, Perry, and Knott counties. We thank the University of Kentucky’s Center for Computational Sciences and Information Technology Services Research Computing for use of the Lipscomb Computing Cluster resources. We are grateful for the laboratory assistance of Nicolette Lawrence, Kara Jones, Mary Foley, and Ricky Grewelle.

## Supplementary Material

**Figure S1:** Mantel correlograms for individual genetic differentiation versus (a) Euclidean distance, (b) natural log of Euclidean distance, (c) stream distance, and (d) natural log of stream distance. Filled circles represent significant values at α = 0.05.

**Figure S2:** Eigenvalue plot (a) and scree plot (b) of local and global axes obtained from our sPCA analyses. The first global axis, in red, was the only axis retained, and displays unique separation from other potential axes in the scree plot (labeled as λ1).

**Figure S3**: Digital elevation model of study area, with sample points corresponding to Figure 1.

**Figure S4**: Visualizations of results from each of the ten replicates for each random subset of loci, and each level of missing data. Includes (a) interpolated vector scores from sPCA, (b) plotted scores from sPCA, and (c) plotted MEM scores for significant results.

**Table S1:** Landcover reclassification scheme for our ResistanceGA resistance surface.

**Table S2**: Summary statistics for read depths from m = 4, m = 5, and m = 7.

**Table S3:** Model output from our ResistanceGA least-cost path analyses. A null model of no geographic structure and a model of isolation-by-distance outperformed all combinations of resistance surfaces based on historic roads, current roads, and surface mining.

