## Supplemental Tables and Figures for "A spatial genomic approach identifies time lags and historic barriers to gene flow in a rapidly fragmenting Appalachian landscape"

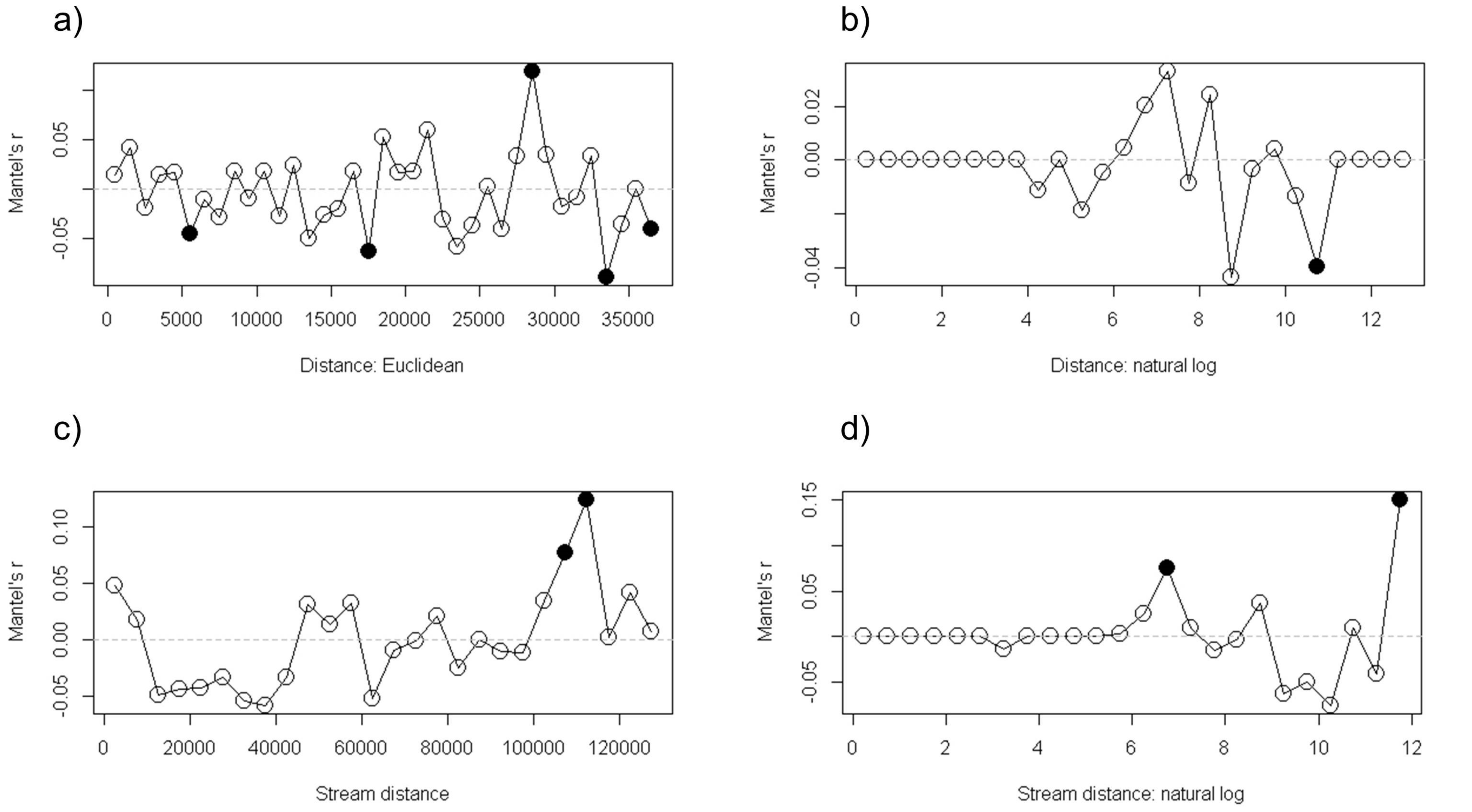

a)
Mantel's r
0.05
-0.05
0
5000
10000
15000
20000
25000
30000
35000
Distance: Euclidean
b)
Mantel's r
0.02
0.00
-0.04
0
2
4
6
8
10
12
Distance: natural log
c)
Mantel's r
0.10
0.05
0.00
-0.05
0
20000
40000
60000
80000
100000
120000
Stream distance
d)
Mantel's r
0.15
0.05
-0.05
0
2
4
6
8
10
12
Stream distance: natural log

a)

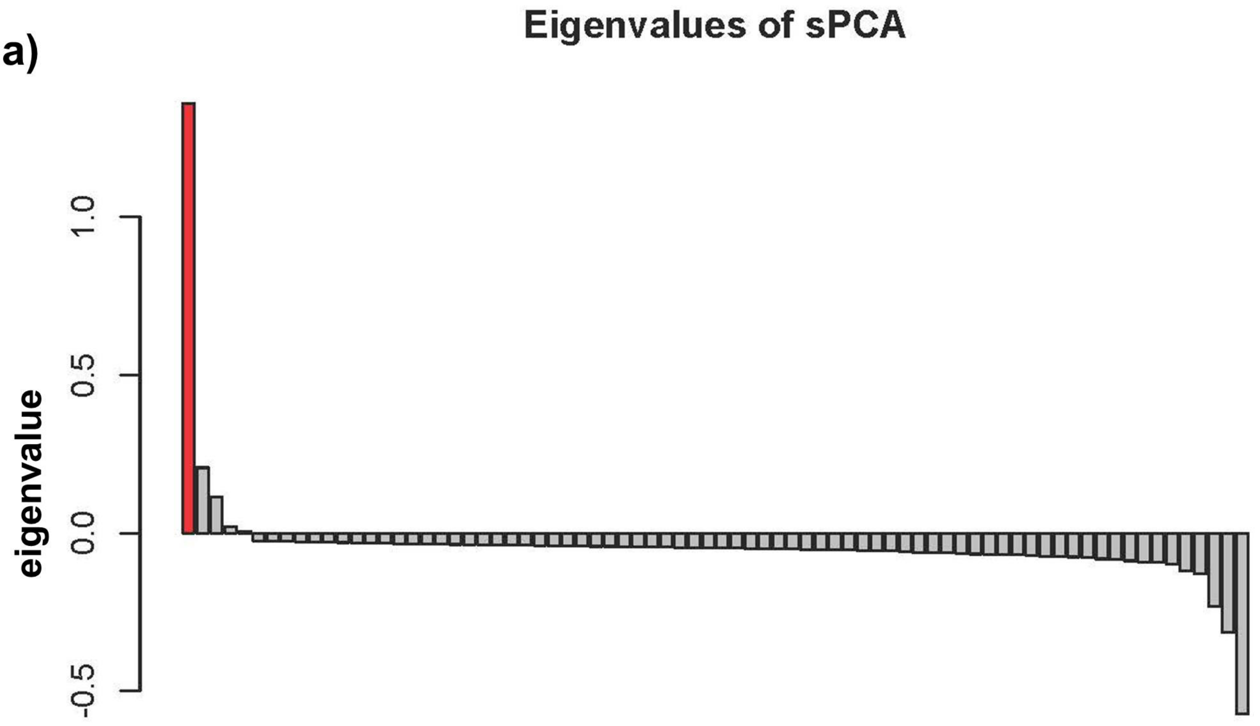
Eigenvalues of sPCA
eigenvalue
1.0
0.5
0.0
-0.5

b)

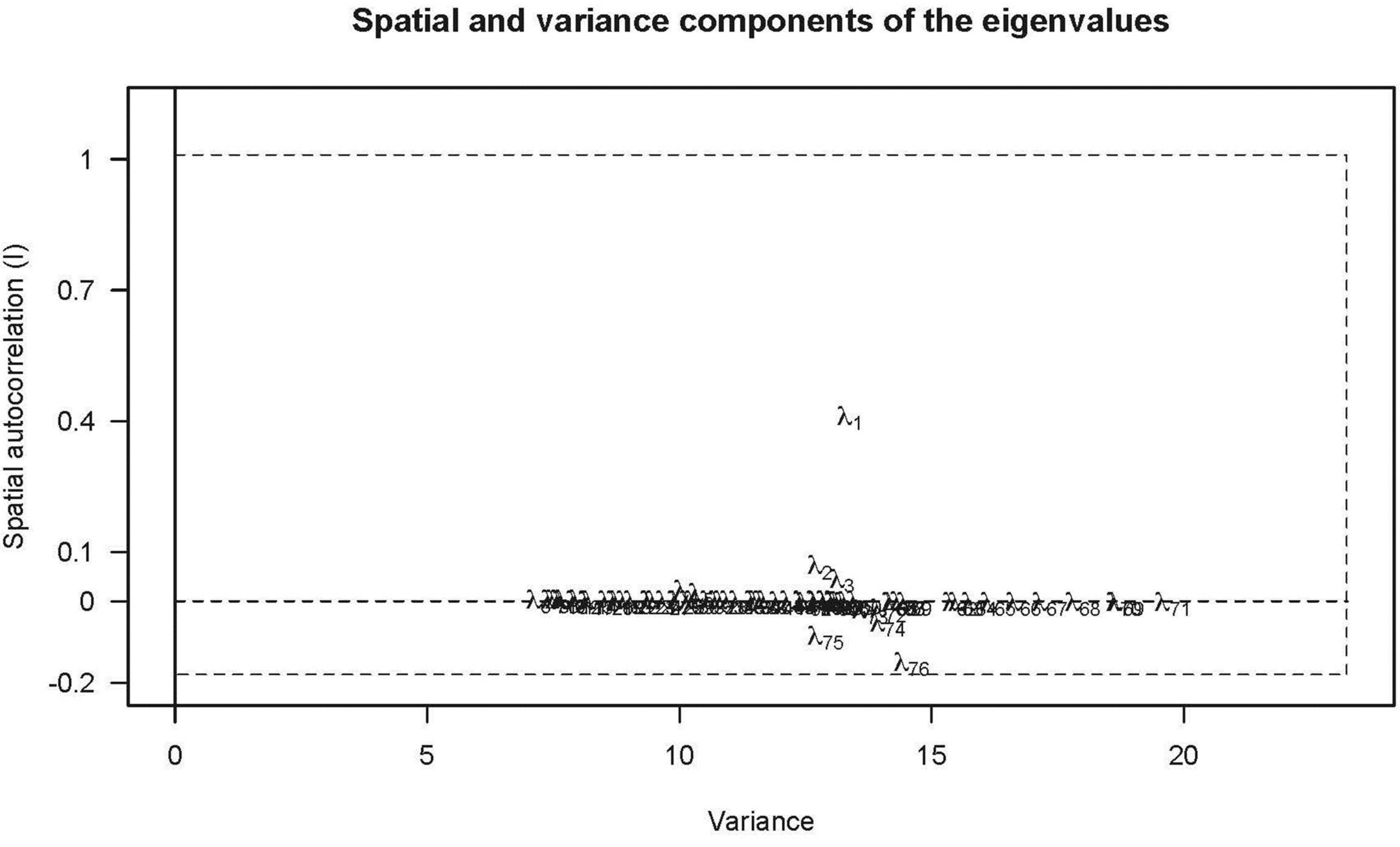
Spatial and variance components of the eigenvalues
Spatial autocorrelation (I)
1
0.7
0.4
0.1
0
-0.2
0
5
10
15
20
Variance
λ1
λ2
λ3
λ74
λ75
λ76
λ71

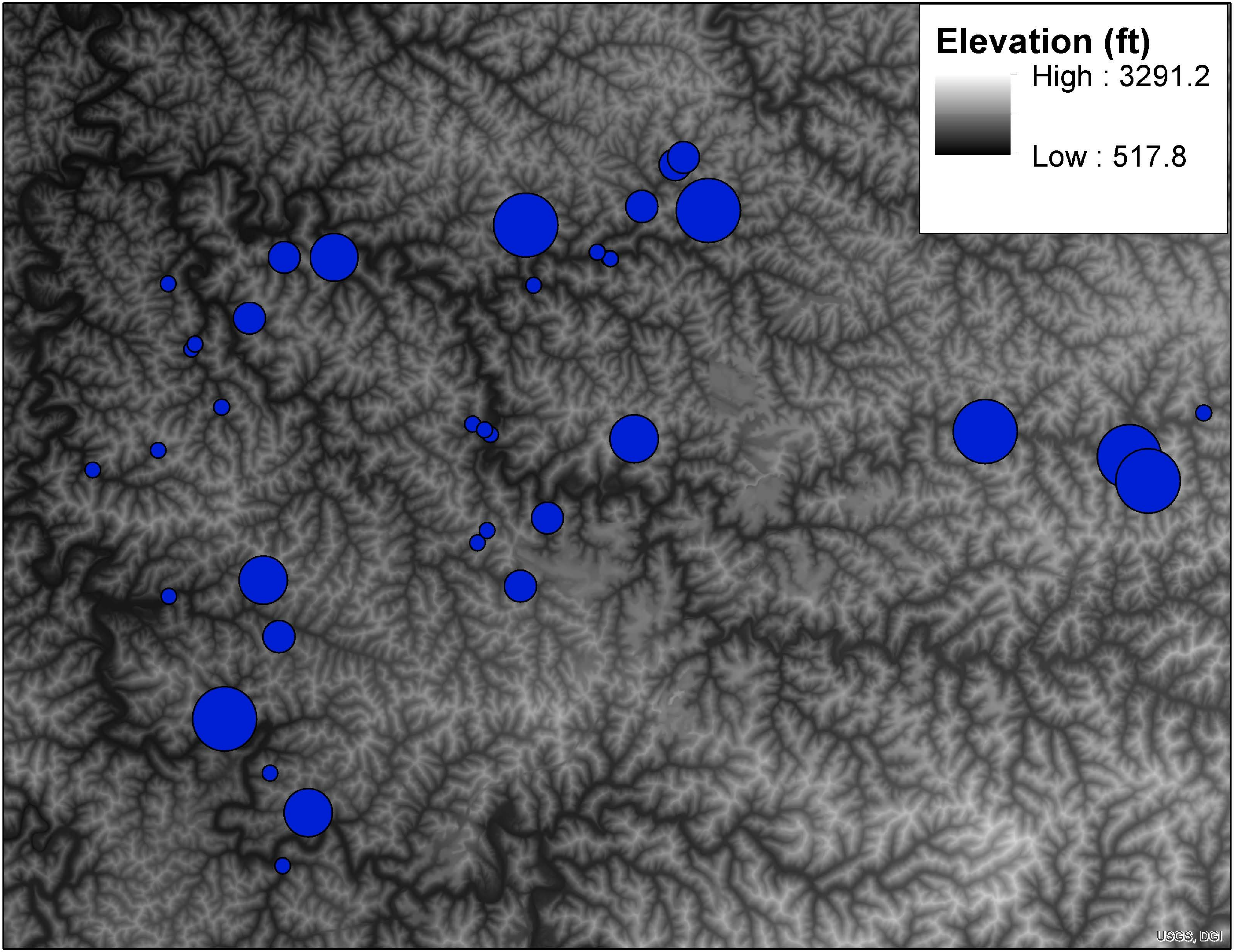
Elevation (ft)
High : 3291.2
Low : 517.8
USGS, DGI

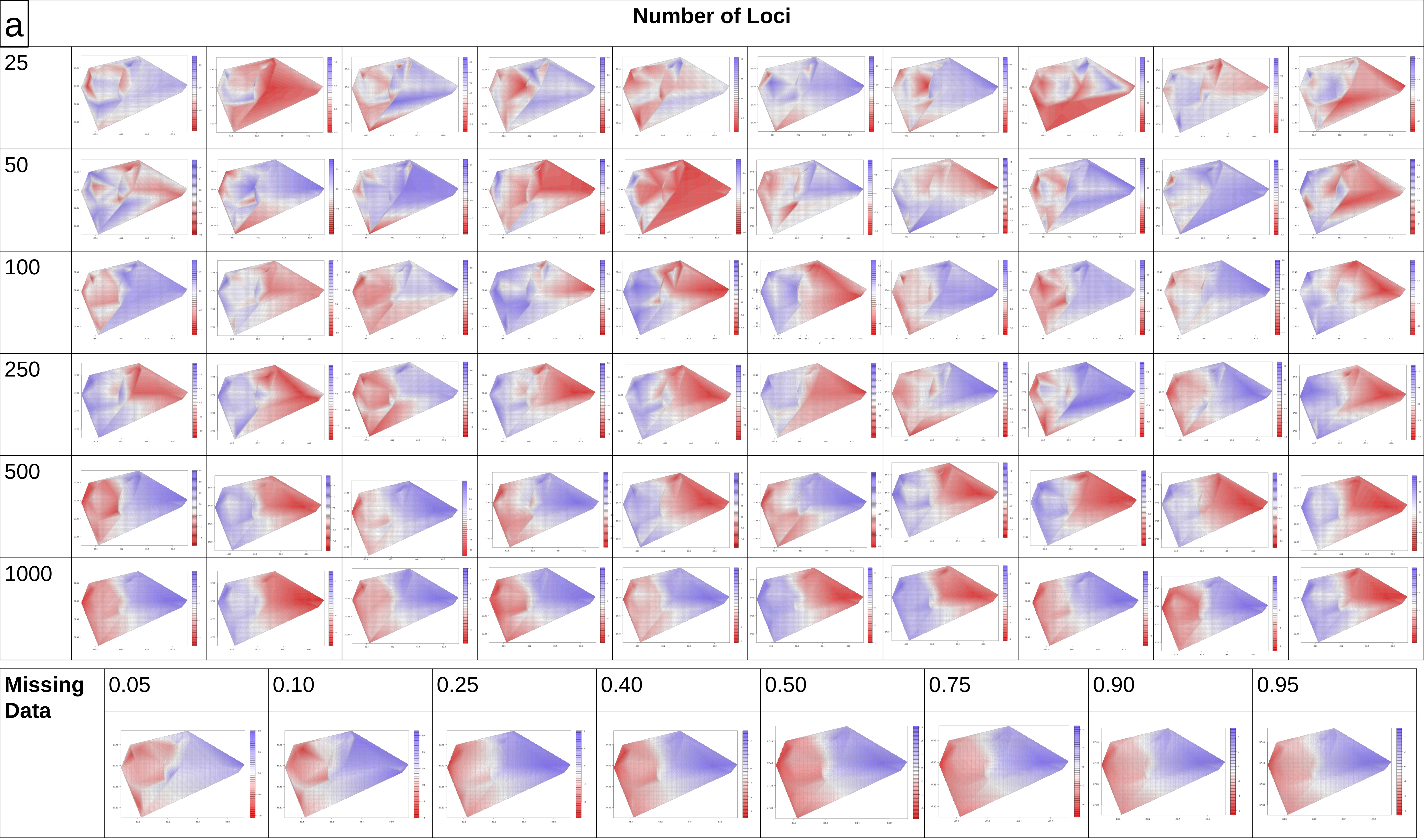
a
Number of Loci
25
50
100
250
500
1000
Missing Data
0.05
0.10
0.25
0.40
0.50
0.75
0.90
0.95

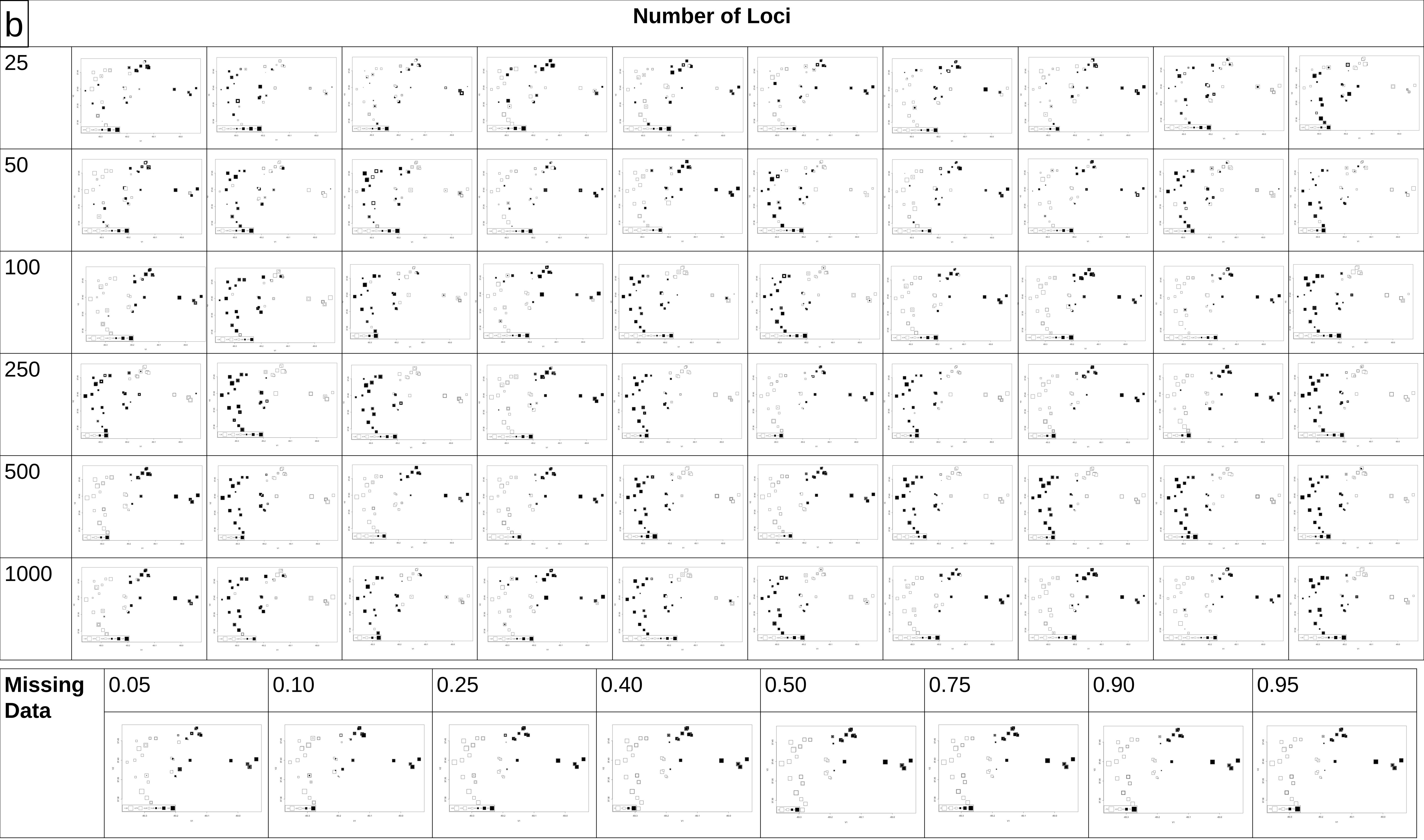
b
Number of Loci
25
50
100
250
500
1000
Missing Data
0.05
0.10
0.25
0.40
0.50
0.75
0.90
0.95

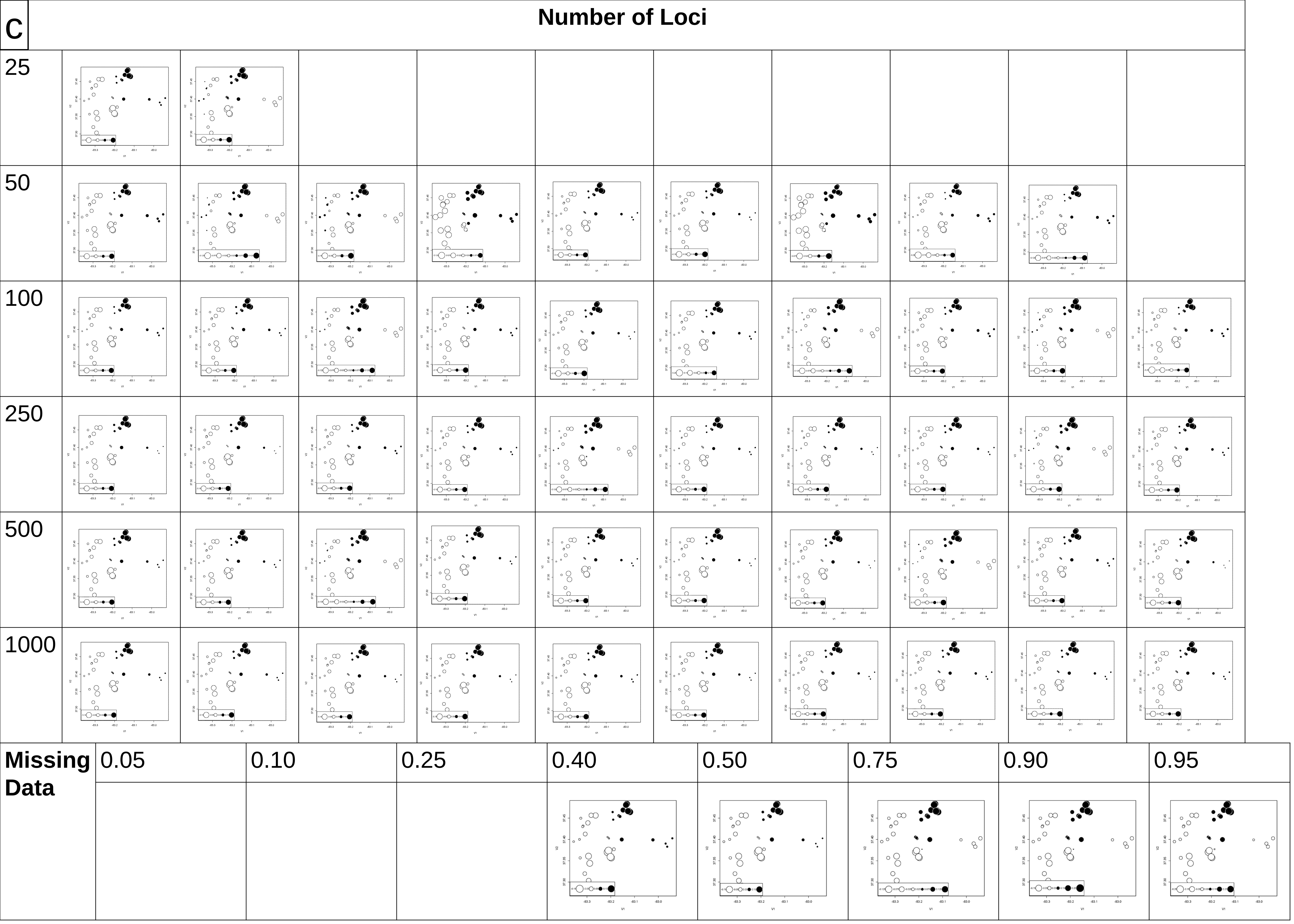
C
Number of Loci
25
50
100
250
500
1000
Missing Data
0.05
0.10
0.25
0.40
0.50
0.75
0.90
0.95

**Table S1:** Landcover reclassification scheme for our ResistanceGA resistance surfaces based on 2011 National Landcover Data classes.

| NLCD 2011 Class | Percent of Study Area | Reclassified Class |
|---|---|---|
| 41 Deciduous Forest | 60.6% | 1 Unmined land |
| 42 Mixed Forest | 0.5% | |
| 43 Evergreen Forest | 4.4% | |
| 90 Woody Wetlands | 0.004% | |
| 95 Emergent Herbaceous Wetlands | 0.005% | |
| 31 Barren Land (Rock, Sand, Clay) | 8.4% | 2 Minelands |
| 52 Shrub/Scrub | 0.1% | |
| 71 Grassland/Herbaceous | 17.5% | |
| 21 Developed, Open | 4.3% | 1 Unmined land[1] |
| 22 Developed, Low Intensity | 2.0% | |
| 23 Developed, Medium Intensity | 0.9% | |
| 24 Developed, High Intensity | 0.2% | |
| 81 Pasture/Hay | 1.0% | |
| 82 Cultivated Crops | 0.3% | |
| Route of KY State Highways 15, 28, 80, and 550 | 0.5% | Current Highways (dichotomous raster, highway/non-highway) |
| Route of current KY 476, northernmost portion of KY 15 | 0.3% | Historic Highways (dichotomous raster, highway/non-highway) |

[1]Where portions of these categories overlap with Reclassified Classes 6 and 7 they are replaced by the reclassified class designation associated with the highway(s).

**Table S2**: Summary statistics for read depths of m = 4, 5, and 7.

| **Read Depth (m)** | **Number of Loci After Filtering** | **Number of Individuals after Filtering** | $H_O$ | $H_E$ | $\pi$ | $F_{IS}$ | $N_e$ **(95% CI)** |
|---|---|---|---|---|---|---|---|
| 4 | 2174 | 77 | 0.199 | 0.244 | 0.245 | 0.184 | 640.9 (600.7, 686.8) |
| 5 | 2140 | 77 | 0.193 | 0.240 | 0.242 | 0.195 | 635.8 (595.6, 681.6) |
| 7 | 2039 | 77 | 0.204 | 0.245 | 0.247 | 0.168 | 629.5 (588.0, 577.1) |

**Table S3:** Model output from ResistanceGA least-cost path analyses. A null model of no geographic structure and a model of isolation-by-distance outperformed all combinations of resistance surfaces based on historic roads, current roads, and surface mining.

| **Surface** | **k** | **AIC** | **AICc** | **$R^2$m** | **$R^2$c** | **LL** | **ΔAICc** | **Weight** |
|---|---|---|---|---|---|---|---|---|
| null | 1 | -10766.7 | -10770.6 | 0 | 0.7057 | 5386.35 | 0 | 0.563 |
| distance | 2 | -10764.8 | -10768.6 | 1.30e-05 | 0.7057 | 5386.39 | 2.034 | 0.204 |
| new_rds | 3 | -10764.9 | -10766.5 | 4.17e-05 | 0.7059 | 5386.44 | 4.110 | 0.072 |
| mines | 3 | -10764.8 | -10766.5 | 2.23e-05 | 0.7057 | 5386.42 | 4.146 | 0.071 |
| old_rds | 3 | -10764.8 | -10766.4 | 1.30e-05 | 0.7057 | 5386.39 | 4.210 | 0.069 |
| mines. + new_rds | 5 | -10764.8 | -10761.9 | 1.30e-05 | 0.7057 | 5386.39 | 8.758 | 0.007 |
| mines. + old_rds | 5 | -10764.8 | -10761.9 | 1.30e-05 | 0.7057 | 5386.39 | 8.758 | 0.007 |
| new_rds. + old_rds | 5 | -10764.8 | -10761.9 | 1.30e-05 | 0.7057 | 5386.39 | 8.758 | 0.007 |
| mines. + new_rds. + old_rds | 7 | -10764.8 | -10757 | 1.30e-05 | 0.7057 | 5386.39 | 13.585 | 0.001 |
